# *I, We*, and *They*: A Linguistic and Narrative Exploration of the Authorship Process

**DOI:** 10.1101/2021.06.14.448236

**Authors:** Abigail Konopasky, Bridget C. O’Brien, Anthony R. Artino, Erik W. Driessen, Christopher J. Watling, Lauren A. Maggio

## Abstract

**Introduction:** While authorship plays a powerful role in the academy, research indicates many authors engage in questionable practices like honorary authorship. This suggests that authorship may be a *contested space* where individuals must exercise *agency*--a dynamic and emergent process, embedded in context--to negotiate potentially conflicting norms among published criteria, disciplines, and informal practices. This study explores how authors narrate their own and others’ agency in making authorship decisions.

**Method:** We conducted a mixed-methods analysis of 24 first authors’ accounts of authorship decisions on a recent multi-author paper. Authors included 14 females and 10 males in health professions education (HPE) from U.S. and Canadian institutions (10 assistant, 6 associate, and 8 full professors). Analysis took place in three phases: (1) linguistic analysis of grammatical structures shown to be associated with agency (coding for main clause subjects and verb types); (2) narrative analysis to create a “moral” and “title” for each account; and (3) integration of (1) and (2).

**Results:** Participants narrated other authors most frequently as main clause subjects (*n* = 191), then themselves (*I*; *n* = 151), inanimate nouns (*it, the paper*; *n* = 146), and author team (*we*; *n* = 105). Three broad types of agency were narrated: *distributed* (*n* = 15 participants), focusing on how resources and work were spread across team members; *individual* (*n* = 6), focusing on the first author’s action; and *collaborative* (*n* = 3), focusing on group actions. These three types of agency contained four sub-types, e.g., supported, contested, task-based, negotiated.

**Discussion:** This study highlights the complex and emergent nature of agency narrated by authors when making authorship decisions. Published criteria offer us starting point--the *stated rules* of the authorship game; this paper offers us a next step--the *enacted* and *narrated approach* to the game.

---

Authorship plays a powerful role in the academy, for individual scholars (e.g., contributing to hiring, promotion, and well-being), for the production of scholarship (e.g., determining which research will be funded and/or published)^1^, and for institutions (e.g., having a reputation for employing highly cited faculty).^2–5^ The International Committee of Medical Journal Editors (ICMJE) has proposed four criteria for authorship, *all* of which must be met for someone to be an author (rather than mentioned in the acknowledgements, for instance).^6^ Yet, while these criteria are helpful for retrospectively attributing authorship, they offer minimal guidance on the prospective *planning* of authorship as one begins a paper and no guidance on authorship *order* once authorship criteria have been met.^7^ Thus, while authors often include ICMJE statements in their papers, a growing body of work indicates that not all authors on published papers actually *meet* the criteria. For example, many survey respondents across disciplines (38% in Rajasekaran et al.,^8^ 52% in O’Brien et al.,^9^ and 62% in Artino et al.’s medical education-specific study^10^) report either participating in or at least observing the practice of “gift” or “honorary authorship” (putting someone on the author line who has not contributed significantly to the work).^11–16^ These studies have also identified factors beyond ICMJE criteria that may influence authorship, including: reciprocity (e.g., to return a favor to someone who helped in some way in the past); feelings of loyalty or obligation, particularly to advisors or mentors; beliefs that the existing criteria do not take into account the important “minutiae of research beyond writing and data reduction”;^15^ and institutional and social structures and hierarchies like tenure status, gender, and minority status.^2,11,15,17–21^ Despite guiding criteria, then, the social practice of authorship remains a *contested* one in that published criteria and enacted practice do not always align.

According to social practice theory, a central mechanism through which social practices are both perpetuated and changed is *agency*, construed as a process mediated by individuals’ social environments and the symbolic resources in those environments.^22^ This process of agency is driven by individuals’ “capacity for understanding and deliberative reasoning that we humans use to select, frame, choose, and execute intentional behavior in the world.”.^23^ This process of agency is dynamic and emergent, and embedded in local and cultural contexts.^23^ Spaces like academic authorship where individuals engage in this process are--by virtue of the personal, local, cultural, and social abutting each other--often contested ones.^22^ In *contested spaces*, individuals must use the cultural and social resources at their disposal to improvise novel ways to solve problems and to exercise agency.^22^ We know little about how authors improvise to negotiate potentially conflicting norms from the ICMJE, their institutions and disciplines, and informal authorship practices.

This complex and often improvisational process of agency can be mediated through language and, in particular, narrative.^23–26^ Through narrative, individuals weave together symbolic resources (i.e., language and gesture) to negotiate an identity within and across social and cultural contexts, co-constructing their own and others’ agency amidst contested spaces.^22^ Narratives can paint a rich portrait of the agency process, using grammatical structure to construct the self, others, and even objects or ideas as acting upon, being acted upon, experiencing, having, and being.^23–26^ Narrating agency, then, is not simply about narrating the self as acting, but it is about narrating self and others in a web of actions, experiences, and relationships.

Despite the growing research on authorship practices, we still understand little about how authors navigate and narrate the complex process of authorship. ICMJE guidelines narrate a straightforward set of rules, but the emergent literature suggests that there is more to the story. In a time of increased collaboration, interdisciplinarity, and team science--where the authorship practices of different individuals and fields are colliding--it is important that we better understand how authors narrate their own and others’ agency in their contexts of authorship. This will allow us to make our practices more visible so that we can prevent injustice and inequity.

In this study we asked: how do authors narrate their own and others’ agency in the process of publishing a paper?

## Methods

We approached this mixed methods study using linguistic and narrative analysis from a pragmatic orientation.^27^ As part of a larger study, we interviewed first authors of recent multi-author papers in medical education and asked open-ended questions about authorship conversations; this analysis focused on participants’ answers to a question about authorship decisions on a recent multi-author (at least three authors) paper (see Maggio et al., 2019, for further details on data collection).^20^ After excluding two participants for incomplete data for this question, our sample included 24 (14 female, 10 male) researchers from U.S. and Canadian institutions, representing assistant (*n* = 10), associate (*n* = 6), and full (*n* = 8) professors. This study was declared exempt by the Uniformed Services University’s Institutional Review Board (protocol #HU-MED-83-9684).

### Data Analysis

We defined as an “account” participants’ verbatim transcribed responses to the question, “Did you have an authorship conversation for this paper and, if so, can you walk me through it?”

These accounts ranged from 164 to 710 words in length (*m* = 382, *SD* = 166). We conducted this mixed-methods analysis in three phases. In phase one, drawing from previous work on how grammatical structure can reflect agency,^25,26,28–30^ AK and LAM independently identified the subjects and predicates of each main clause (a subject and a tensed predicate without subordinating conjunctions) in the account and categorized these subjects as singular first author *I*, first-person plural *we* (first author and others), other authors (either singularly--*he, she*--or in a group--*they*), non-author persons (e.g., editors), or inanimate (either specific nouns like *the discussion* or *the paper* or nonspecific pronouns like *there* or *it*). Following Halliday and Matthiessen’s functional linguistic schema, AK and LAM also categorized predicates as material (verbs of doing like *circulate, put, work on*), mental (experience verbs like *think, know, feel*), verbal (communication verbs like *offer, ask, talk*), or relational (verbs establishing a relationship like *have* or *be*) types of actions.^31^ Finally, AK and LAM came together to resolve disagreements, coming to consensus on all linguistic coding.

In phase two, AK, LAM, and BCO examined each account as a narrative, focusing on the “work” participants were doing with these stories.^32^ Two researchers read each story and, using the participants’ words as much as possible, gave each one a title (conceived as what the story was “about”) and a moral (conceived as what the point of the story was and based in part on Labov and Waletzky’s notion of a coda).^33^ Each pair met to compare their titles and morals and, reviewing evidence from each story, came to consensus.

In phase three we integrated the results of phases one and two, creating a profile for all participants that included (a) the most frequent main clause subject type, (b) the most frequent predicate type, and (c) the title and moral of their authorship story. Balancing the quantitative data of *who* participants narrated doing *which* actions (i.e., subjects and predicates) with the qualitative data of the topic and purpose of the story (i.e., titles and morals), AK interpretatively characterized each participant narrative into a type of agency. LAM and BCO reviewed these interpretations to determine if they fit with their own sense of the data. The entire team then met to discuss the agency type assigned to each profile and participated in shaping their interpretations.

### Reflexivity

As authors of a piece on authorship, we felt it important to explore how we experienced the authorship process. All authors reflected in writing and then orally as a group on what we perceived ourselves and others bringing to the project, our expectations for authorship and author order, and how this research may influence how we approached authorship for this manuscript. AK analyzed written responses and notes from the oral reflection after completing analysis of participant data. See the appendix for the results of this reflexivity exercise.

## Results

As Table 1 shows, across all subjects and ranks, other authors were the most frequent subjects (e.g., *they, she, he*), followed by first-person *I*, then inanimate nouns and pronouns (e.g., *the paper, it*), then *we*. Participants narrated relational verbs the most frequently followed by material and, less often, mental and verbal. Tables 2 and 3 show the distribution of subject and process types by gender and rank, respectively.

**Table 1.** Total Subject and Process Types Across Participants

|  | **Coded Instances** |
| --- | --- |
| I | 151 |
| We | 105 |
| Other author | 191 |
| Inanimate | 146 |
| Non-author animate | 7 |
| Material | 193 |
| Mental | 90 |
| Relational | 241 |
| Verbal | 76 |
| Total coded sentences | 600 |

**Table 2.** Mean Number of Subjects and Process Types by Gender (N = 24)

|  | **F**<br>**mean (SD)** | **M**<br>**mean (SD)** |
| --- | --- | --- |
| I | 5.9 (6.2) | 6.9 (3.3) |
| We | 5.1 (3.5) | 3.3 (3.5) |
| Other author | 7.7 (6.6) | 8.3 (7.4) |
| Inanimate | 5.5 (3.2) | 6.9 (4.7) |
| Non-author animate | 2 (.3) | 2 (.3) |
| Material | 9.3 (7.4) | 6.3 (4) |
| Mental | 3.4 (2.5) | 4.3 (2.3) |
| Relational | 9.4 (3.4) | 11 (6.6) |
| Verbal | 2.5 (1.8) | 4.1 (2.7) |
| Total coded sentences | 24.5 (12) | 25.7 (9.6) |

**Table 3.** Mean Number of Subjects and Process Types by Rank (N= 24)

|  | **Assistant**<br>**mean (SD)** | **Associate**<br>**mean (SD)** | **Full**<br>**mean (SD)** |
| --- | --- | --- | --- |
| I | 7.8 (6.5) | 6 (3.3) | 4.6 (4.3) |
| We | 4.9 (3.8) | 5.3 (4.1) | 3 (2.9) |
| Other author | 6.9 (3.5) | 8.7 (9.6) | 8.9 (8.2) |
| Inanimate | 6.5 (2.8) | 5.8 (2.5) | 5.8 (5.8) |
| Non-author animate | 2 (.2) | .7 (.8) | 1.3 (.4) |
| Material | 8.6 (7.5) | 9.2 (7.7) | 6.5 (2.8) |
| Mental | 4 (2.4) | 4.7 (2.2) | 2.8 (2.5) |
| Relational | 10.2 (4.9) | 9 (3.6) | 10.6 (6.3) |
| Verbal | 3.5 (2) | 3.7 (1.2) | 2.4 (3.2) |
| Total coded sentences | 26.3 (11.4) | 26.5 (12) | 22.3 (10.3) |

Based on the interpretive integrating process in phase three, we found three broad types of agency with which authors narrated the authorship process: (1) *distributed*: either (a) other author and inanimate or non-referential nouns (e.g., *it, there*) subjects with material (e.g., *work, support*) or verbal (e.g., *discuss*) actions or (b) *we* subjects with relational states (e.g., *be, have*); all with a narrative focus on cataloging the contributions from across the team, (2) *individual*: *I* subjects with material actions or relational states; narrative focus on what the self as first author brought to the project, and (3) *collaborative*: *we* subjects with material or verbal actions; narrative focus on the joint actions of the team. Below we describe each of these types and the subtypes (supported, contested, task-based, and negotiated) that emerged in this data set. (See Table 4 for a summary.)

**Table 4.** Agency Types and Sub-Types from this Sample

| **Agency Type** | **Agency Sub-Type** | **Linguistic Features** | **Description** | **Number of Participants** |
| --- | --- | --- | --- | --- |
| Distributed | Supported | -relational verbs<br>-*we* or other author subjects | agency of the team critically connected to resources distributed across the team and supported by authorship norms | 10 |
|  | Contested | -relational verbs<br>-*it* subjects | resources and norms of team contrasted with strong individual agency assertion | 2 |
|  | Task-based | -material verbs<br>-other author subjects | separate material actions of each team member drive the process | 2 |
|  | Negotiated | -verbal verbs<br>-other author subjects | verbal agency of team members drives the authorship process | 1 |
| Individual | Supported | -material or relational verbs<br>-*I* subjects | self's actions and resources are primary drivers, but are supported by actions and resources of the team | 6 |
| Collaborative | Negotiated | -verbal with some material & relational verbs<br>-*we* subjects | verbal actions along with material actions and resources of team drive authorship process | 2 |
|  | Contested | -material verbs<br>-*we* subjects | material actions and resources and norms of teams contrasted with strong individual agency assertion | 1 |

**Table 5.** Exemplar Participants for Each Broad Agency Narrative Type

|  | **Participant D**<br>(Female, Associate) | **Participant I**<br>(Female, Assistant) | **Participant U**<br>(Male, Assistant) |
| --- | --- | --- | --- |
| Agency type | *Distributed supported* | *Individual supported* | *Collaborative negotiated* |
| Most frequent subject | other authors (45%, $n = 13$) | I (42%, $n = 23$) | we (53%, $n = 8$) |
| Most frequent verb type | relational (62%, $n = 18$) | material (51%, $n = 28$) | material and verbal (33% each, $n = 5$) |
| Narrative features | resources of the team (i.e., what they *are/have*) drive the process | self's material actions drive the process with support from other authors | material and verbal actions of the team drive the process |
| Title | Breaking with "the older paradigm" for the senior author and recognizing "who's doing the most work" | "I...circulated it [manuscript] around": Coordinating a multi-site paper | "It got turned down...they made us go back and get more": Adding authors after an initial submission |
| Moral | "The two people that are doing the most work should be first and last"—there is a new paradigm among junior faculty | "I didn't have a big conversation with everybody, but...this is a pretty cordial group...and there were no issues that I was aware of anyway"—with a cordial group, all goes well | "It [authorship] wasn't an issue"—with "conversation," everyone can end up happy |
| Brief summary | The account begins with Participant U recounting how each author is connected to a data set, then moving through not just what each “does” but who they “are” (e.g., educator, potential coding tiebreaker). | The account begins with a group at a conference writing a proposal. Her actions coordinating the effort are supported by what “we” did and the absence of barriers (expressed via relational verbs) like “difficulties.” | The account begins with a smaller author team getting an article rejected. Participant D narrates how “we” brought on more authors, making their middle authorship clear “up front.” |

### Distributed Agency

This was the most common type of agency, narrated by 15 participants. In distributed agency, the most frequent subjects were the other authors (i.e., not the participant--sometimes including *we*), inanimate (e.g., *the decision, the paper*), or non-referential (e.g., *there, it*) nouns; the most frequent process types were relational, material, or verbal; and the titles and morals, while varying a bit, tended to focus on the contributions, expectations, or roles of various members of the author team. Because there were so many examples of this type of agency, we were able to parse out four subtypes based on different combinations of subjects, process types, and morals, which we describe below.

### Distributed supported agency

Ten participants narrated distributed supported agency, characterized by frequent relational verbs and *we* or other authors as subjects. These authors narrated a largely conflict-free experience, with morals focusing on how either existing relationships or patterns of work across the project made authorship roles “clear,” “easy,” “comfortable,” “understood,” or other phrases reflecting ease. All of these stories relied on relational verbs to narrate who the team *was* or what resources the team *had* that allowed them to succeed. Often these resources were in the form of relationships, an author who “was my thesis advisor” (Participant Y) or a group of authors who were “people I’ve worked with on multiple projects” (Participant P). Each author was narrated as bringing a different set of skills, knowledge, experiences, and relationships to the table to support the overall project.

### Distributed contested agency

The two participants who narrated with this type of agency, like those in the category above, relied on relational verbs to narrate the team’s resources. In contrast, however, these participants also emphatically narrated their own *individual* contributions, contesting the broader distributed narration and defending their own position as first author. Participant R said, “I wrote it, full stop,” referring to the process as “blood, sweat and tears” and Participant Z that it was “clear” she would be first author because she did “the lion’s share of the work.” Yet neither author narrated team conflict and both focused their narratives on others’ contributions. In fact, the former narrated how the team presented a “united front” on a potentially controversial issue and the latter referred to the “wonderful colleagues” on her team. Rather than contestation as conflict, then, it is about emphasizing the *individual* contributions amidst the distributed process.

### Distributed task-based agency

The two participants using this agency type also focused on the role of other authors, but they narrated those roles more as material than relational; they talked more frequently about what individual authors *did* rather than what they *were* or *had*. Both morals were about the relationship between work and authorship, with Participant S saying, “we kind of took the approach that the more you did, the further to the outside [of the authorship list] you were.” Participant A’s moral problematized this work/authorship relationship, referring to a more senior team member who was perhaps more involved in the conceptual than the writing process: “whether or not he typed the words, he was definitely a part of the writing.”

### Distributed negotiated agency

One participant’s distributed agency account focused on the *verbal* actions of himself and the author team. As with most other participants in this category, Participant T’s most frequent subjects were other authors, but his focus was less on their resources or what they do and more on what they say, agree upon, or have a conversation about. This distributed negotiated agency led, in Participant T’s view, to the whole process “end[ing] up being pretty simple because it was all done up front.”

### Individual Supported Agency

Six participants narrated what we might think of as a more typical type of agency, with *I* subjects being most frequent. These participants narrated themselves *working, feeling, thinking, updating* and *circulating* drafts, *deciding* on methodological approaches, *asking* other authors questions, and *doing* the IRB, among other processes. Yet their narratives all demonstrated, as Participant G puts it, that these authors--for better or for worse--were all “pretty securely embedded in the author group.” Thus, while they may have narrated the self’s actions more frequently, they also narrated this agency being *supported* by the team’s actions and resources. For instance, they talked about how other authors *clarified* concepts, *were* “senior people” (Participant G) respected in their field, *were* “old and good friends” (Participant K) and *took on* an active role. They also narrated collaborating, like *we met* and *put together* a proposal, *started hashing out* ideas, *circulated* material, *drafted up* the paper, *decided* on author order, and *divided up* the work.

### Collaborative Agency

The remaining three participants told stories of collaborative agency, in which *we* was the most frequent subject, the most frequent verbs were material or verbal, and titles or morals pointed to collaborative experiences or conversations. Two of these participants told a *collaborative negotiated* story with morals suggesting that a “conversation up front” (Participant M) laying out expectations created a positive working environment. These participants narrated a process void of team conflict and peppered their stories with verbal *we* actions like *discussing, talking about* and material ones like *bringing on* (another author) and *sitting down* (to compose an outline). Participant F, however, told what we call a *collaborative contested* story, noting both that “it was very much a collaborative experience” and that “I definitely wrote the entire paper myself.” This story begins and ends with individual *I* actions, sandwiching the collaborative *we* actions and, to some extent, contesting the *we* actions.

## Discussion

The participants in this study narrate three broad types of authorship agency--distributed, individual, and collaborative--with four possible subtypes: supported, contested, task-based, and negotiated. These narratives highlight the complex and emergent nature of agency:^23^ for instance, some participants who narrated distributed agency (*we*, other author, and *it* subjects most frequent) focused more on the *relationships* among authors and resources (i.e., contested and supported agency, where relational verbs are most frequent) while others focused more on each author’s *material* actions (i.e., task-based agency, where material verbs are most frequent) and one focused most on what authors’ *said* (i.e., negotiated agency, where verbal verbs are most frequent). Additionally, these narratives do not seem to reflect the collaborative approach we might have predicted for this multi-authored research: while distributed and individual agency were relatively common (42% and 25% of narratives respectively), collaborative agency was relatively infrequent (13% of narratives). These results suggest that authorship is a nuanced set of practices with a variety of distinct approaches and understandings.

Yet the seven sub-types of agency identified in these narratives represent only a portion of the *possible* approaches and understandings. For instance, while we identified only individual *supported* agency (*I* subjects and material or relational verbs most frequent) among these participants, this same combination of subjects and verb types could result in an individual *contested* agency if the participant contested the more frequent individual agency by emphatically noting at one point the importance of what *we* collaboratively did. Similarly, an author could narrate an individual *negotiated* agency with *I* subjects and verbal verbs. This potential variety of understandings arose from the profile we created for each narrative, drawing from both linguistic (frequency of subjects and verb types) and narrative (titles and morals) analysis. This integrated approach was particularly helpful in understanding the agency in the sometimes shifting and uncertain context of authorship teams, and we maintain it could be applied to other “messy” problems to more fully understand agency in contested spaces like interprofessional collaboration, remediation, and trainee shame experiences.^34–36^

While our methods captured some of the messiness of authorship practices, these narratives did not reflect the conflicted nature of authorship that the literature on responsible conduct of research might suggest.^2,9–15,17–21^ Participants did not narrate overt conflict related to these 24 papers (only three narratives were contested, and these contested elements were subtle) and frequently stressed the helpful actions and resources of team members. Rather than reflecting the improvisational negotiation of conflicting norms,^22^ these narratives suggested that authorship is an agreeable space of productive work. Optimistically, this could mean that authorship practices have evolved to feature explicit conversations, both early and throughout the experience, that mitigate later feelings of regret, resentment or discomfort. It might also represent our sample of first authors (who may be less aware of conflict or inequity elsewhere in the author list) with published papers (who, therefore, are happy overall with the process). Yet, few participants brought up authorship criteria, so the absence of conflict might mask an acceptance of certain practices that may not fit guidelines but that contribute to keeping everyone happy (e.g., including a senior person even though they did no writing). In this sense, it is possible that few participants had their agency tested, because they felt comfortable with the cultural norms that shaped the decision making. Alternatively, they may have felt they could not contradict these cultural norms and, hence, did not raise any issues. Further research might probe for conflict in authorship practices, as well as examine agency in non-first authors, to further explore the cultural norms of authorship.

The absence of conflict in these narratives may also signal the potential for inequity in authorship practices. Racist practices, policies, and cultural norms are a major source of inequity in medicine and other health professions fields.^37–44^ This is indicative of what Okun calls white supremacy culture, one aspect of which is fear of conflict and an accompanying belief that those in power have a right to feel comfortable.^44^ Institutions like medicine tend to use politeness and silence around the “uncomfortable” topics of race and racism as two tactics to maintain white supremacy.^44^ In order to dismantle white supremacy culture, Okun argues, institutions must learn to “distinguish between being polite and raising hard issues.”^45^ Equitable assignment of authorship is indeed a hard issue and is something only few of the participants narrate addressing overtly with team members. Our approach breaks authorship practice down into the people and resources at play (the subjects), the actions, experiences, and relationships of the authorship team (the verb types), and the overarching beliefs (in the form of morals and titles) team members have about the process. This offers scholars a vocabulary of sorts to begin to discuss conflict and the “hard issues” of racial--and other--inequities in authorship.

### Limitations

There are several limitations to this study. First, we only interviewed first authors of published papers, omitting the perspectives of the rest of the author team and, further, scholars who were unsuccessful in getting published. This may put a more rosy perspective on authorship than the full range of stories would. Second, while we discuss the absence of conflict in the narratives, we did not probe participants about potential conflict. While we expected that participants would describe the full authorship experience, including the conflict, future work could directly examine conflict. Third, we did not ask participants’ about their race; future studies should explore the authorship experiences of those underrepresented in healthcare.

Finally, our sample was drawn from authors based in the United States and Canada. It is possible, had we sampled more broadly, we would have observed different narratives.

## Conclusion

To our knowledge, this study is the first to explore how HPE authors narrate their own and others’ agency in the authorship process. Taking agency to be a complex process, embedded in social and cultural contexts, we were able to discern different *types* of narratives authors use to make sense of their experiences. The ICMJE criteria offer us a starting point--the *stated rules* of the authorship game; these data offer us a next step--the *enacted* and *narrated approach* to the game. Distributed, individual, and collaborative narrative types offer another starting point not just for our conversations about authorship, but for our conversations *about* those conversations: how do we want to talk about the work we do and the relationships we form as we do them?

## Acknowledgments

We would like to thank Joe Costello for all his help and support with the piece.

## Appendix A

### Results of Reflexivity Exercise

We found that our written feedback reflected more individual (asking what “I” think “I” bring to the project) and distributed (asking what “others” contribute) agency and our oral feedback more collaborative (how are “we” making decisions and referring to “our work).

Authors’ written reflections narrated what each saw the self (e.g., “I think I also bring qualitative expertise” [BCO]) and specific others (e.g., “Abby brings unique methodological expertise and contributes more than most of the other authors” [ED]) bringing to the table. Meanwhile, the group oral reflection centered more on who “we” are and what “we’’ bring, usually moving away from this particular project towards comparison with past projects and with a focus on the importance of the relationships and the work versus authorship. All concurred that the mostly senior composition of the team makes authorship “less contentious” in the words of AA. AA also notes that he would feel “uncomfortable” asking for “higher billing” and others agree. In this group context, then, we lean towards narrating collaborative agency.

Finally, towards the end of our conversation, AA noted in a humorous aside that gender most likely played a role in past negative author experiences. Our mixed gender group (in which women have the coveted first, second, and last author spots) responded with laughter and the topic died. Reflecting on this later over email, AK, CW, and LM noted the complexity of factors at play such as female team members’ methodological and content expertise, male team members’ sensitivity to implicit bias and privilege, issues of academic identity and whether authors see this work as part of their central “academic canon” [CW], and indeed the reflexivity session itself.

